# Direct control of shell regeneration by the mantle tissue in the pearl oyster *Pinctada fucata* via accelerating CaCO3 nucleation

**DOI:** 10.1101/572024

**Authors:** Jingliang Huang, Yangjia Liu, Taifeng Jiang, Wentao Dong, Guilian Zheng, Liping Xie, Rongqing Zhang

## Abstract

Molluscan bivalves rapidly repair the damaged shells to prevent further injury. However, it remains unclear how this process is precisely controlled. In this study, we applied scanning electronic microscopy, transmission electronic microscopy and histochemical analysis to examine the detailed shell regeneration process of the pearl oyster *Pinctada fucata*. It was found that the shell damage caused the mantle tissue to retract, which resulted in dislocation of the mantle zones to their correspondingly secreted shell layers. However, the secretory repertoires of the different mantle zones remained unchanged. As a result, the dislocation of the mantle tissue dramatically affected the shell morphology, and the unusual presence of the submarginal zone on the nacreous layers caused de novo precipitation of prismatic layers on the nacreous layers. Real-time PCR revealed that the expression of the shell matrix proteins (SMPs) were significantly upregulated, which was confirmed by the thermal gravimetric analysis (TGA) of the newly formed shell. The increased matrix secretion accelerated CaCO_3_ nucleation thus promoting shell deposition. Taken together, our study revealed the close relationship between the physiological activities of the mantle tissue and the morphological change of the regenerated shells.

## 1. Introduction

Organisms are capable of depositing a diverse array of minerals, which fulfill important biological functions. One of such functions is to protect the body from predator attack. Accordingly, the predators strengthen their weapons (teeth and claws). The arms race between the predators and the preys drives the evolution of remarkable skills for survival and results in the extraordinary biominerals with outstanding mechanical properties, such as limpet teeth (Mann et al., 1986), sea urchin spines (Seto et al., 2012), crustacean exoskeletons (Chen et al., 2008; Raabe et al., 2005), and molluscan shells (Song et al., 2003). Among these, molluscan shells have been extensively studied due to their hardness and toughness, which make them as ideal models for bioinspired ceramics (Finnemore et al., 2012; Jackson et al., 1988).

The molluscan shells can be rapidly repaired when external aggressions occur, which endows the molluscs undeniable evolutionary advantage. Shell regeneration induced by artificial damage is widely adopted to reveal the shell formation process, because the regenerated shells resembled the normal shells and the repair process was similar to normal shell deposition (Chen et al., 2019; Huning et al., 2016b; Meenakshi et al., 1974). Usually, shell regeneration begins with deposition of an organic membrane (Chen et al., 2019; Pan and Watabe, 1989), serving as the temporary barrier and the first substrate for the mineral phase deposition. Although the shell regeneration is conducted by the shell secreting mantle, the morphology of the repaired shells may slightly differ from the normal shells, as found in the green ormer *Haliotis tuberculate* (Fleury et al., 2008) and the mussel *Mytilus edulis* (V. R. Meenakshi, 1973). Such discrepancy may due to the stress response of the mantle tissue. Indeed, our recent study showed that, in the pearl oyster *Pinctada fucata*, Peroxidase-like protein and β-N-acetylhexosaminidase were exclusively expressed during the shell repair process (Chen et al., 2019) and might be involved in the initiation of the prismatic layer formation. However, it remains unknown how the mantle tissue response to the shell damage stimulation and how its physiological changes affect the shell morphology.

The pearl oyster *P. fucata* have been extensively studied in the biomineralization field. The shell of *P. fucata* consists of inner nacreous layers and outer prismatic layers. The nacreous layers are hundreds of layers of aragonitic tablets separated by organic matrix, resembling the brick-mortar walls. The prismatic layers contain dozens of layers of longitudinally-arranged columnar calcite. Each prismatic layer is coated by a periostracum membrane on the outer surface. The formation of the shell has been ascribed to the matrix secreting mantle tissue (Marie et al., 2012; Zhang and Zhang, 2006). The mantle tissue can be divided into three regions according to their different secretory repertoires: mantle edge, submarginal zone, pallial and central zones (Fang et al., 2008). The mantle edge is responsible for the periostracum formation and initial stage of the prismatic layer deposition (Suzuki, 2013). The submarginal zone further thickens the prismatic layer, while the nacreous layers are secreted by the pallial and central zones of the mantle tissue (Marie et al., 2012). The shell formation process is precisely controlled by the mantle tissue.

In this study, we are seeking to understand the whole process of the shell regeneration after shell damage. Scanning electron microscopy (SEM), transmission electronic microscopy (TEM), thermal gravimetric analysis (TGA), histrochemical analysis and real-time PCR were used to examine both the regenerated shells and the covering mantle tissue. The results showed that the nucleation of CaCO_3_ was promoted by upregulating the SMPs secretion in the mantle tissue.

## 2. Materials and methods

### 2.1 Oyster collection and cultivation

The pearl oyster *Pincatada fucata* was obtained from Guangdong Ocean University (Zhangjiang, China) and air transported to Beijing. The oysters were acclimated for one week in an aquarium tank containing 700 L artificial sea water (salinity 33.0 ± 0.5 psu, pH 8.1 ± 0.05) at room temperature. The oysters were fed twice a week with commercial Spirulina before and during the experiment.

### 2.2 Artificial shell damage-induced shell regeneration

Totally 80 healthy individuals with dorsal-ventral shell length of 6-7 cm were randomly selected for experiment. A “V” nick on the shell was made by cutting the ventral edge with a scissors (Figure 1). The cut shell pieces were examined to make sure that the inner nacreous layers were injured. The oysters were then returned to the tank and collected at 6 hours (h), 12 h, 24 h, 48 h, 7 days (d), 30 d and 60 d after the treatment. At each time point, six individuals were anaesthetized by soaking the oysters in 1000 mL sea water containing 0.25 % phenoxy propyl alcohol for 10 min. Six untreated oysters were used as a control group. Then the oysters were fixed with 4 % formaldehyde in sea water for 24 h.

**Figure 1.**
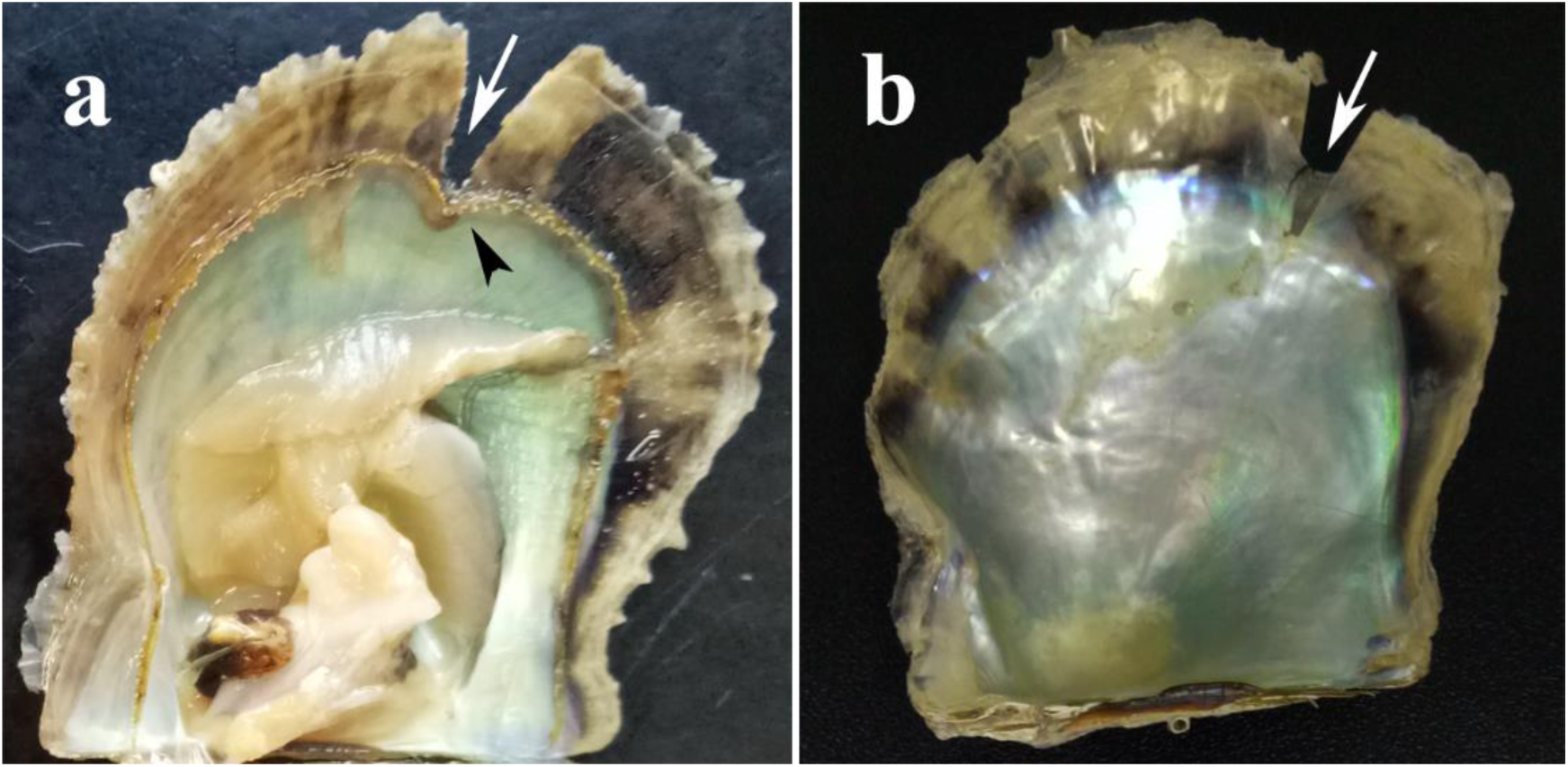
Artificial shell damage in *Pinctada fucata*. a, formaldehyde-fixed oyster sample with the left valve removed, 48 hours after shell damage. Note that the mantle tissue retracted into the pallial zone (black arrow head) at the notching site (white arrow). b, shell sample of 30 days after shell damage showing the thin regenerated shell layer covering the notching site (white arrow).

### 2.3 Scanning electron microscopy (SEM) analysis

After removing the covering mantle tissue, the shell samples containing the “V” nick and the adjacent area were cut by a scissors and a glass cutter. The small shell pieces were coated with gold and examined by a scanning electron microscope (SEM, FEI Quanta 200, Germany) with an accelerating voltage of 30 kV in a high vacuum mode.

### 2.4 Decalcification of the shell and histochemical analysis

The regenerated shells were cut by a scissors after removing the mantle tissue and used for subsequent decalcification. Specifically, we prepared the mantle-shell sample with mantle remained attaching to the shell inner surface. To obtain such samples, the adductor muscle was cut by a scalper after the fixation mentioned above, and then the mantle covering the injury was carefully separated from the gill and the adjacent mantle region by a razor blade not to make any displacement of the mantle-shell. The inblock was cut by a scissors and a glass cutter. All the shell samples were completely decalcified with 0.5 M EDTA and rinsed three time in sterile water. The decalcified samples were paraffin-embedded after a gradient ethanol dehydration. A routine histochemical procedure of H&E stain was subsequently applied and an Olympus IX81 light microscope was use to photograph the slices.

### 2.5 Thermal gravimetric analysis (TGA)

The regenerated shells from 30 oysters of 60 days after shell damage were collected by nipper and merged. These samples were mainly prismatic layers. As a control, prismatic shell layers from 10 normal individuals were collected and merged. The shell samples were ultrasonic washing in ddH_2_O three times and air dried. The content of the organic compounds in the shells was measured by TGA (TherMax, Cahn Instruments, China) in a nitrogen atmosphere. The heating temperature ranged from room temperature to 900 °C at a rate of 10 °C per min.

### 2.6 RNA extraction

To separately extract RNA from the mantle edge and pallial zone, the treated and untreated oysters were immersed in one litter sea water containing 0.25 % phenoxy propyl alcohol for 10 min. After the animal was fully anaesthetized, the adductor muscle was cut by scalpel carefully. Then the edge and pallial zone of the mantle around the notching site were cut and held in 2 ml RNase-free Eppendorf tubes. For each time point, tissues from six animals were merged into one sample. Each sample was added with 1 ml Trizol (Thermo Fisher Scientific, USA) and stored at −80 °C.

For the RNA extraction, two steel balls (pretreated at 180 °C for four hours to denature any RNase) were added to each sample tube after unfreezing the tissues. A tissue breaker (TL2010S, DHS, China) equipped with a high speed shaker was used to grind the mantle tissues, and the homogenized mixture was transferred to a new tube. Then 200 µL chloroform was added to denature the protein components. The mixtures were vortexed and centrifuged at 12000 g 4 °C for 15 min. The supernatants (∼600 µL) were transferred to new tubes and added with 150 µL chloroform. The mixtures were vortexed again and centrifuged at 12000 g 4 °C for 15 min. The supernatants were transferred to new tubes and mixed with isopropyl alcohol of equal volume. The solutions were gently mixed and kept at −20 °C for 10 min. Then a centrifugation (12000 g, 4 °C, 15 min) was applied, and the supernatants were discarded. The RNA pellets were rinsed with 1 ml 75 % alcohol for once and air dried in a clean bench. The RNAs were dissolved in 40 µL RNase-free water. The quality and concentration of the RNAs were examined by Nanodrop 2000 (Thermo Scientific, USA).

### 2.7 Reverse transcription and real-time PCR

PrimeScript™ RT Master Mix (TaKaRa, Shiga, Japan) was used to reverse transcribe the RNA into cDNA. And real-time PCR analysis of the gene expressions were performed according to our previous study (Huang et al., 2018) using SYBR® Premix Ex Taq™ product (TaKaRa, Shiga, Japan) in a StepOnePlus™ Real-Time PCR System (Applied Biosystems, Vernon, CA, USA). Gyceraldehyde-3-phosphate dehydrogenase (GAPDH) was used as an internal reference, and the primers for the shell matrix proteins are listed in Table 1. The relative gene expression levels were calculated by the -ΔΔCT method.

**Table 1.**
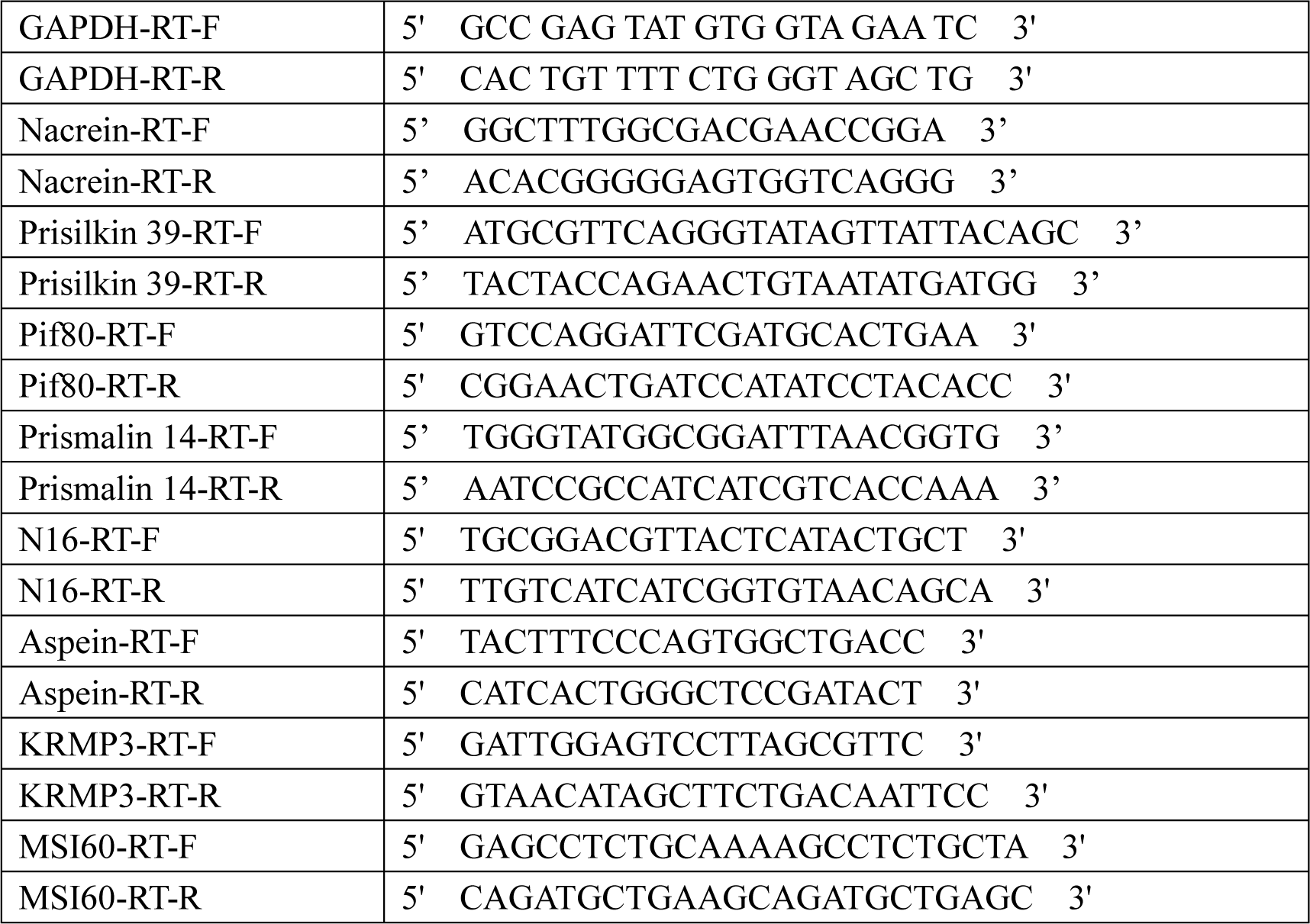
Primers for real-time PCR in this study.

## 3. Results and Discussion

### 3.1 The general shell regeneration process

We performed a long term study of the shell regeneration in the pearl oyster *Pinctada fucata*. No mortality due to the shell damage was observed up to 60 days, and all the oysters exhibited shell regeneration to varied extent. The mantle edge retracted into the pallial zone soon after the notching treatment and remained staying behind the cut edge (Figure 1a). As the repair progress, a transparent shell sheet began to grow right upon the injured site (Figure 1b), which could be seen as early as 7 days, consistent with previous studies (Chen et al., 2019; Huning et al., 2016a), until the nick was progressively covered by newly formed shell layers. The mantle edge surrounding the nick also displayed a “V” shape arrangement, indicating that the outer epithelium is capable to recognize the physical condition of the shell surface, although the manner by which is not clear.

The regeneration was quite rapid. At 6 hours after the shell damage, thin organic membrane was evidenced near the nick (Figure 2a) and supposed to be the periostracum which is the initiation of prismatic layer deposition (Suzuki, 2013). Another important role of the periostracum was to set up a barrier to enclose the extrapallial space from the ambient sea water. The periostracum was continued to be secreted and expanded in the following hours (Figure 2b and 2c). Simultaneously, the microstructure of the adjacent nacreous layer was affected (Figure 2d). The retracted mantle edge and submarginal zone might disturbed the normal nacre deposition. Alternatively, the dissolution of hexagonal aragonitic tablets might be due to the hypoxia (Melzner et al., 2011; Silverman et al., 1983), caused by the close of the shell valves during the first few hours post shell damage. Numerous particles were found within the periostractum at 24 h (Figure 2e) and began to grow as the periostracum continue to be secreted at 48 h (Figure 2f). Suzuki et al. (Suzuki, 2013) showed that the initial growth of the prism column begins with the nucleation of calcium carbonate in the periostracum. Consistently, in the early stage of shell regeneration, periostracum was first laid down on the previous shell layers following by the deposition of calcium carbonate particles which would further grow into prism. The nacreous layers were covered by disordered crystals with a relatively smooth and flat morphology at 48 h (Figure 2g). At 7 d after shell damage, a newly formed shell layer was visible around the nick (Figure 2h) and was found to be prism (Figure 2i). The inner surface of the prism was rough and composed of nanograins, in accordance with our previous study (Chen et al., 2019). At the dorsal side of the regenerated shell layer, an atypical prism/nacre transition zone was observed (Figure 2j). In normal condition, nacreous layers grow and spread upon the inner surface of the mature prismatic layers, as seen in Figure 2a. However, at the early stage of shell regeneration, precipitation of nacre tablets was interrupted at the injury site and replaced by prism deposition. The temporal transition zone was composed of grains of several microns (Figure 2k). As the repair proceeds, the regenerated prism covered and bridged the dorsal part of the nick at 30 d (Figure 1b and Figure 2l). In some individuals, the nicks were completely covered and the regenerated shells were comparable to the shells before damage in length at 60 d. However, the microstructure of the regenerated prismatic layers (RPL) (Figure 2m and 2n) was quite different from the that of the normal ones (Figure 2a and 2o), suggesting that the shell regeneration process is a long time event. As shown in Figure 2n, the RPL layer was deposited right on the periostracum, following by a secondary prism. The primary prism contained prolonged granules of several microns in diameter, which were not well shaped and embedded in the organic matrix. The secondary prism appeared to be more developed with the prism columns were clearly shaped, although the diameters (around 10 microns) were significantly smaller than those of the normal prisms (around 50 microns, Figure 2o).

**Figure 2.**
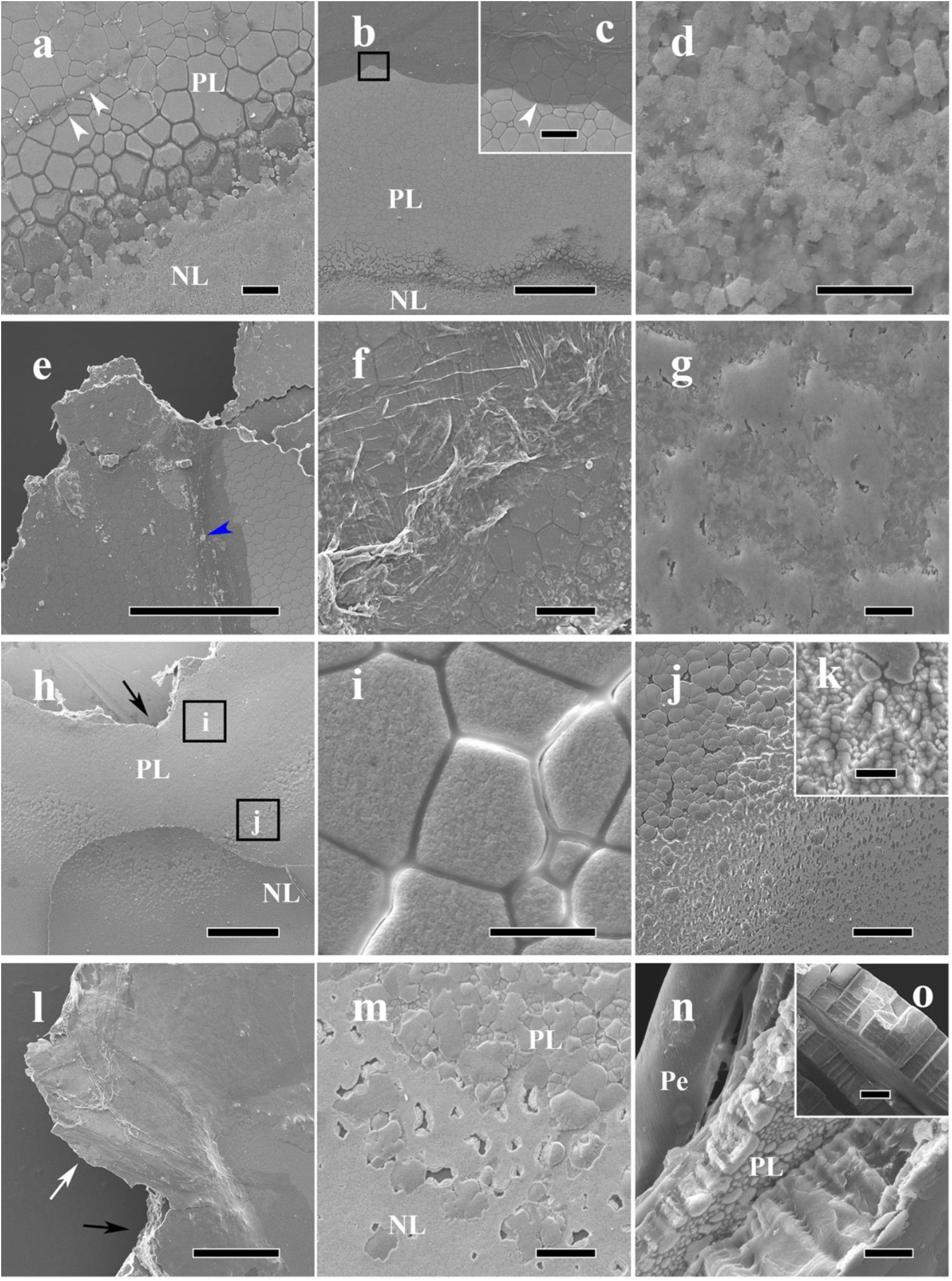
SEM observation of the regenerated shells. a, 6 hours after shell damage, transparent organic membrane (periostracum) was visible near the notching site (white arrow heads). b-d, 12 hours after shell damage, showing that the adjacent prismatic layer was covered by an periostracum membrane (arrow head in c) and the nacreous layer deposition was affected (d). c is the magnification of the black frame in b. e, 24 hours after shell damage, CaCO_3_ depositions were visible within the covering periostracum membrane (blue arrow head). The notching site is at the top left. f and g, 48 hours after shell damage. The periostracum membrane was thickened and more particles were deposited in the membrane in the adjacent prismatic layer (f) and nacre tablets were no longer visible in the adjacent nacreous layer (g). h-k, 7 days after shell damage. i and j are the magnifications of the two black frames in h, showing the newly formed prism polygons (i) near the notching site (black arrow) and the atypical prism/nacre transition zone (j). k is the magnification of j. l, 30 days after shell damage, the previous prismatic layer (black arrow) was covered by a thin regenerated prismatic layer (white arrow). m and n, 60 days after shell damage, showing the recovering prismatic and nacreous layers. n is the side view of the regenerated prism layers. o, side view of normal prism layers. PL, prismatic layer; NL, nacreous layer; Pe, periostracum. Scale bars in b, e, h and l are 500 µm; scale bars in a, c, f, j, m and o are 50 µm; scale bars in d, g, i, k and n are 10 µm.

To further study the regenerated shells in detail, we decalcified the shell samples and performed H&E staining. As shown in Figure 3, after removing the CaCO_3_ by EDTA, the remaining matrix frameworks of the nacre were blue-violet in color, while those of the prism was purplish red. Interestingly, the framework of the regenerated prismatic layers slightly differed from the normal prism and were in dark red, indicating that their compositions might not be exactly the same. Another feature revealed by the histochemical analysis was the peculiar Sandwich structure of the shell layers, which is consistent with Figure 2h. This phenomenon could be clearly figured out in Figure 3h and 3j. In such situations, the general prism-nacre depositing order has been reversed, in other words, the RPL were deposited on the previous nacre and followed by regenerated nacreous layers (RNL). The RPL deposition began with the formation of mature periostracum which was seen at 48 h after shell damage (Figure 3b and 3f) but not in samples of 12 h (Figure 3a and 3e). The slightly differences between the SEM observation and the H&E stain might be ascribed to the high resolution of the SEM.

**Figure 3.**
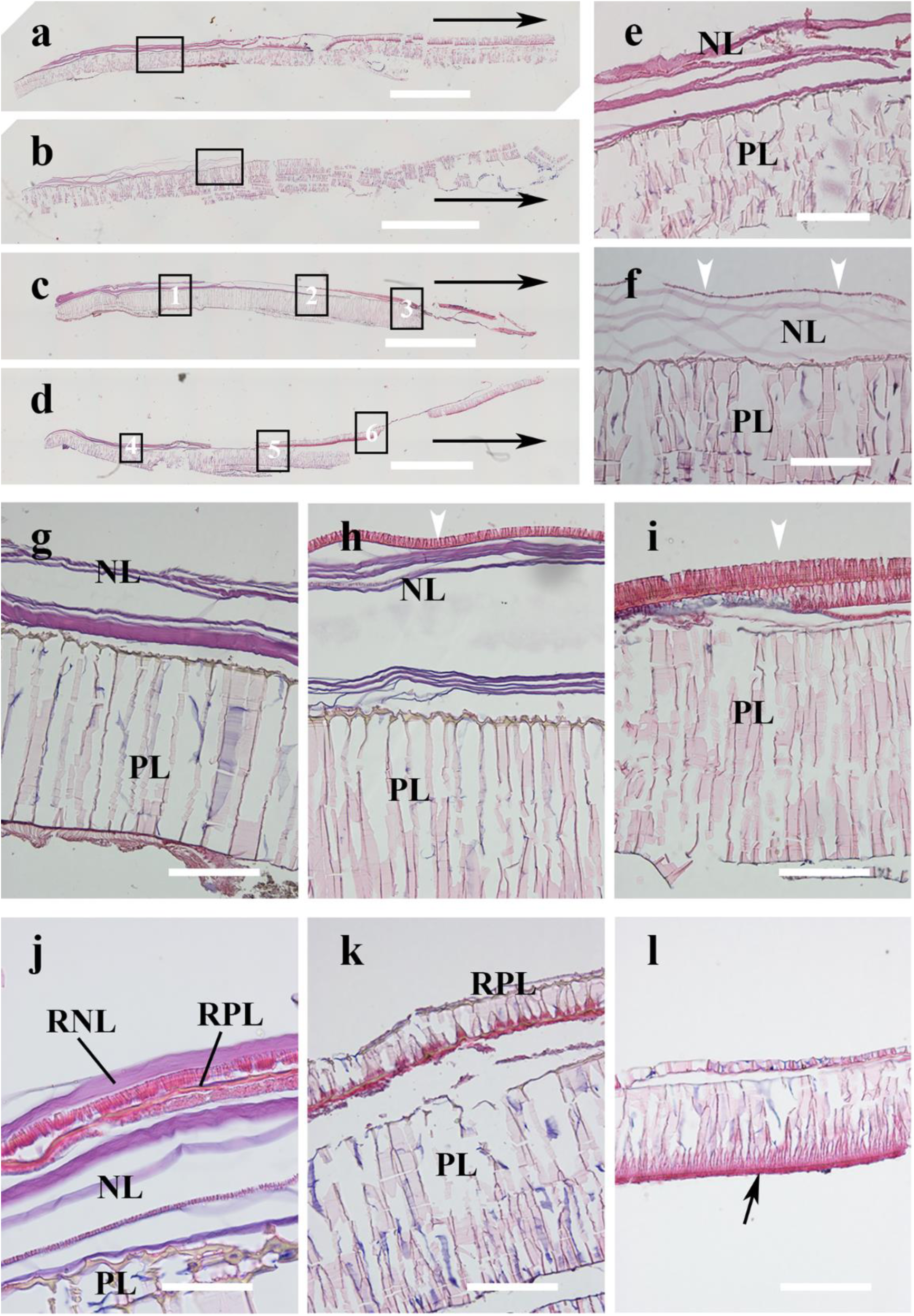
H&E stain of the decalcified shell samples after shell damage. a-d, panorama view of the decalcified shells of 12 hours, 48 hours, 7 days and 30 days after shell damage, respectively. The long black arrows in a-d indicate the growth direction of the shells. e and f are magnifications of the black frames in a and b, respectively. The white arrow heads indicate the newly formed periostracum. g-i are magnifications of c, corresponding to black frames 1-3, respectively. The white arrow heads in h and i indicate the regenerated prismatic layers. j-l are magnifications of d, corresponding to black frames 4-6, respectively. The black arrow in l indicates the periostracum membrane. PL, prismatic layer; NL, nacreous layer; RPL, regenerated prismatic layer; RNL, regenerated nacreous layer. Scale bars in a-d are 1mm; the others are 100 µm except j (50 µm).

### 3.2 The nucleation sites of primary prismatic layer

As observed in the histochemical analysis, the primary layers of the RPL were composed of tiny prisms compared with the large prism columns formed in normal conditions (Figure 3h and 3i), suggesting that nucleation of calcium carbonate in the RPL was dramatically promoted. Because each prism column can be regarded as one nucleation event of calcium carbonate in the initiation stage of prismatic layer formation (Ubukata, 2001). Indeed, when we looked into the outer surface of the regenerated prismatic layers which represent the initial stage of the shell repair process, the morphology was quite different. At day 7 after shell damage, the primary layer contained intensive irregular prisms which were crowded and in tower shape (Figure 4b and 4d). The size was 5-15 µm in diameter, much smaller than the normal prisms (Figure 4a) which were 30-50 µm in diameter. The number of the tower prisms was around 63.6 per 10^4^ µm^2^, much more than the normal prisms (17.1 per 10^4^ µm^2^ on average). At day 30 after shell damage, the number of prisms was 21.5 in 10^4^ µm^2^ similar to the normal ones, indicating the nucleation rate of CaCO_3_ fell down to basal line (Figure 4e). This was further confirmed by the SEM result (Figure 4c). However, the periostracum at day 30 after shell damage was thicker than the control, and abundant organic materials filled between the prism columns. Moreover, the diameters of the prisms were dispersed, indicating the asynchrony of the nucleation events. These results showed that the shell repair process is an emergency response with accelerated mineralization.

**Figure 4.**
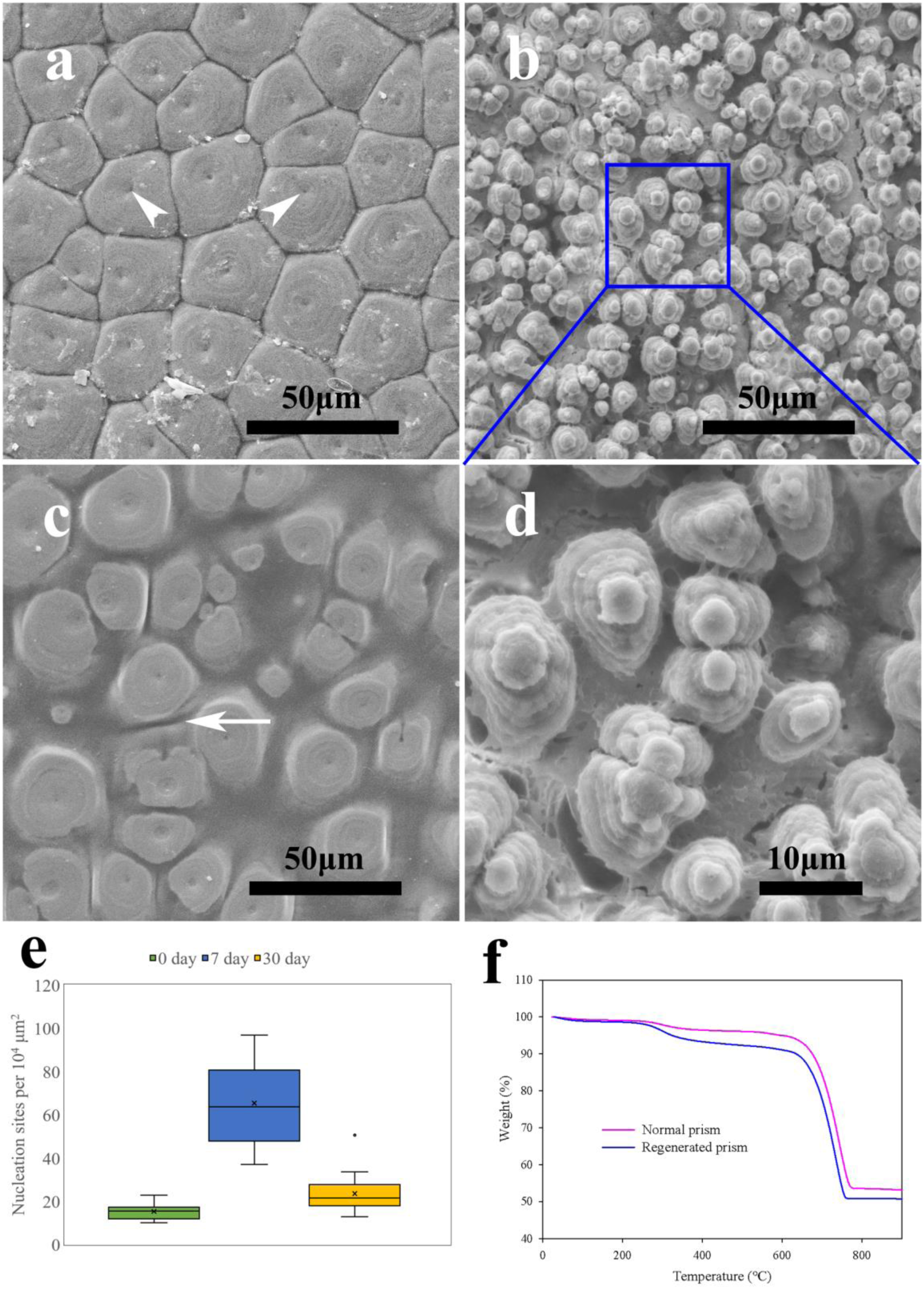
a-d, SEM images of the outer surface of the prismatic layers, representing the nucleation events of CaCO_3_. a, normal prismatic layer with regular polygons. White arrow heads indicate the nucleation sites within each prism. b and d, outer surface of a regenerated prismatic layer at day 7 with the periostracum layer peeled off. d is the magnification of b. Note that the initial small prisms are in tower shape. c, outer surface of a regenerated prismatic layer at day 30. The white arrow indicates the organic material between the prisms. e, quantitative analysis of the initial nucleation events of prismatic layer during the shell regeneration process (n=6; in each shell sample, 2-3 areas were examined). f, TGA analysis of normal and regenerated prismatic layers (30 days after shell damage).

The prismatic layers in bivalve shells contain high content of organic compounds which play vital roles in the shell formation. We found that organic materials in the prism of *P. fucata* was about 4.05% of the bulk weight, consistent with those of other bivalves (2.7-6.1%) (Checa et al., 2005). The organic materials include matrix proteins, polysaccharides, lipids and other small molecules. And the matrix proteins are proved to be involved in calcium carbonate nucleation, polymorphs selection, crystal orientation, and have been extensively studied (Liang et al., 2015; Miyamoto et al., 1996; Ponce and Evans, 2011; Takeuchi et al., 2008). It has been demonstrated that many matrix protein genes were upregulated after shell damage stimulation (Kong et al., 2009; Lin et al., 2014) and some SMPs have been proved to promote nucleation of calcium carbonate, such as PfY2 (Yan et al., 2017), Alv (Kong et al., 2018) and Prismalin-14 (Suzuki et al., 2004). Therefore, we speculate that the mantle tissue promotes the nucleation of calcium carbonate by secreting more organic matrix, thus accelerating the shell regeneration process. Indeed, we found that the regenerated prism contained more organic materials than normal prism layer (Figure 4f). The weight losses between 230°C and 600°C were due to the thermolysis of the organic matter, and the release of CO_2_ from the decomposition of CaCO_3_ after 600°C led to further dramatic weight losses (Li et al., 2017). The content of the shell matrix in the regenerated prismatic layers was calculated to be about 7.46% weight of the total mass, probably the highest matrix content in biominerals.

### 3.3 Direct control of the mantle tissue on the shell repair process

Mantle tissue plays a central role in the shell formation. To understand how the mantle conducts the shell regeneration, special mantle-shell samples were prepared (Figure 5a and 5e). The anesthesia treatment before fixation resulted in the well preserved morphology of the tissues close to their physiological state. It was found that the mantle was in direct charge of the regeneration process. Right at the injury site, the mantle retracted to the nacre region (Figure 1a). In this manner, the ventral part of the nacre was covered by the mantle edge and submarginal zone which secreted a regenerated prismatic layer upon the former (Figure 5a-5d). It follows that the displacement does not alter the secretary repertoires of the mantle edge and submarginal zone (see the following section). As the growing tip of the shell was propelled forwards, the mantle gradually repositioned. As a result, the RPL upon the previous nacre would be covered by the homing pallial zone of the mantle, and the latter would deposit layers of regenerated nacre upon the RPL (Figure 3j).

**Figure 5.**
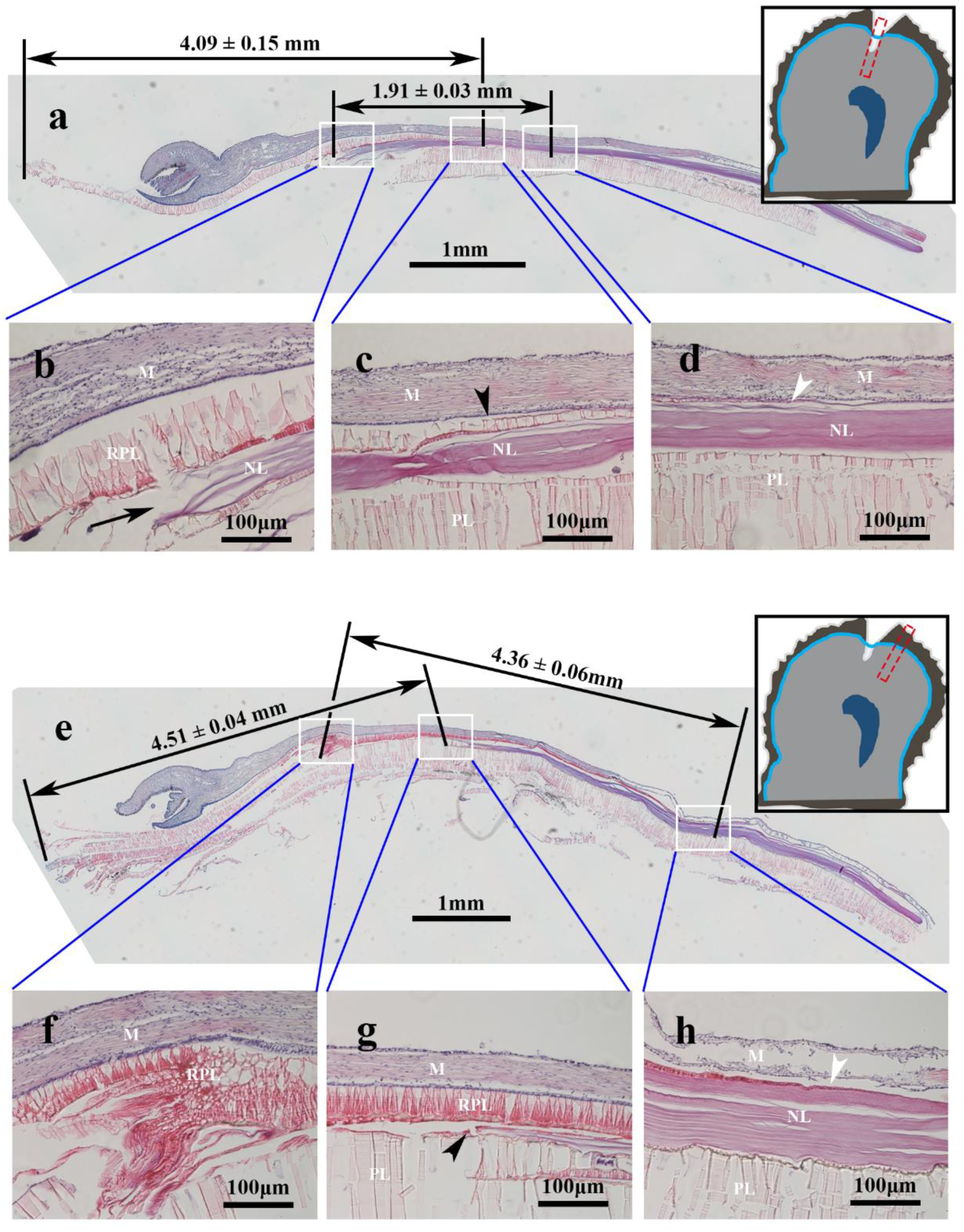
H&E stain of the mantle-shell samples at 60 days after shell damage. a, decalcified shell sample with the mantle tissue covering the notching site, as indicated by the inset at the top right corner. b-d, the magnification of the white frames in a, showing the cut edge (black arrow), forming prism/nacre transition zone (black arrow head) and the starting position of the regenerated prismatic layer (white arrow head). e, decalcified shell sample with the covering mantle tissue, taken from the adjacent region paralleled to the notching site (as indicated by the inset on the right). f-h, the magnification of the white frames in e, showing the retraction line of the mantle edge, previous prism/nacre transition zone (black arrow head) and the starting position of the regenerated prismatic layer (white arrow head). Note that the corresponding lengths were measured along the silhouette of the shells.

As shown in Figure 5, the morphology of the regenerated shells and the behavior of the mantle were closed related. Right at the notching site, the area between the cut edge of shell notching (Figure 5b) and inner-most of the RPL (Figure 5d), was measured about 1.91 ± 0.03 mm in length, corresponding to the retracted mantle edge and submarginal zone at the very beginning of the shell regeneration process. This length is less than a half of that between the regenerated shell edge and the frontier of the regenerated nacre (Figure 5c), corresponding to the growing prismatic layer, was about 4.09 ± 0.15 mm. Therefore, the shell damage not only caused the retraction of the mantle tissue, but also led to the contraction of the mantle edge and the submarginal zone at the injury site, which affecting the shell morphology in return. However, in the adjacent area parallel to the notching site, the length of the growing prism before shell damage (4.51 ± 0.04 mm) and RPL right at the beginning of shell repair (4.36 ± 0.06 mm) were comparable, indicating slightly contraction of the mantle tissue. Interestingly, on the adjacent inner shell surface, parallel to the nick, accumulation of periostracum was observed (Figure 5f), indicating that the shell damage led to a stationary state of the mantle tissue and the mantle edge kept secreting periostracum without precise control.

Shell matrix proteins (SMPs) fulfill vital roles in CaCO_3_ nucleation, orientation, and crystal polymorph selection (Miyamoto et al., 1996; Ponce and Evans, 2011; Sudo et al., 1997; Takeuchi et al., 2008). And the biomineralization processes of the prism and nacre are controlled by the different SMPs secretary repertoires of the different mantle zones (Marie et al., 2012). Several identified SMPs have been showed to be significantly upregulated after shell notching treatment (Fang et al., 2012; Pan et al., 2014). However, in these studies, the gene expressions were detected in the whole mantle tissue, therefore, it remains unclear whether the SMPs expressions in each mantle zone are precisely controlled. We separately detected the secretory regimes in the mantle edge and the pallial zone, corresponding to the prismatic layer and nacreous layer secretion, respectively. The results showed that post shell notching treatment, the prism SMPs, namely KRMP, Prismalin-14 and Prisilkin-39 were significantly upregulated in the mantle edge (Figure 6a), while the nacre SMPs, namely Pif80, N16 and MSI60 were significantly upregulated in the pallial zone (Figure 6b). Nacrein is present in both prism and nacre, and its expression was upregulated in both the mantle edge and the pallial zone (Figure 6a and 6b). Interestingly, neither prism SMPs were detected in the pallial zone, nor the nacre proteins in the mantle edge during the shell repair process (data not shown). Therefore, the shell damage stimulated the SMPs upregulation in the corresponding mantle zones, but did not change the functional secretory regimes. In a recent study, Anne K. Hüning et al. (Huning et al., 2016a) showed that several genes that are specifically expressed in pallial and marginal zones could be induced in central mantle after experimental injury in the central part of the shell. The shell morphology during the flat pearl formation in the abalone *Heliotis rufescens* (Fritz et al., 1994) and pearl oyster *P. fucata* (Xiang et al., 2013) also suggested that the secretory regime of central part of the mantle tissue are programmable. Such inconsistency may due to the different approaches to induce shell damage, which result in varied shell repair strategies of the molluscs. When shell damage occurred in the central part of the shell, no retraction of the mantle tissues was observed, and the shell repair was accomplished by the central mantle, which might force the central mantle to reprogram the secretory regime to fulfill the deposition of outer shell layers (Fritz et al., 1994). However, in our study, the damage was occurred at the edge of the shell and forced the mantle to retract. Although the mantle edge might sense the unusual signal of the nacre surface, its secretory regime remained unchanged. As a result, the mantle edge deposited a regenerated prismatic layer on the underlying nacre and determine the re-initiation of the shell formation process.

**Figure 6.**
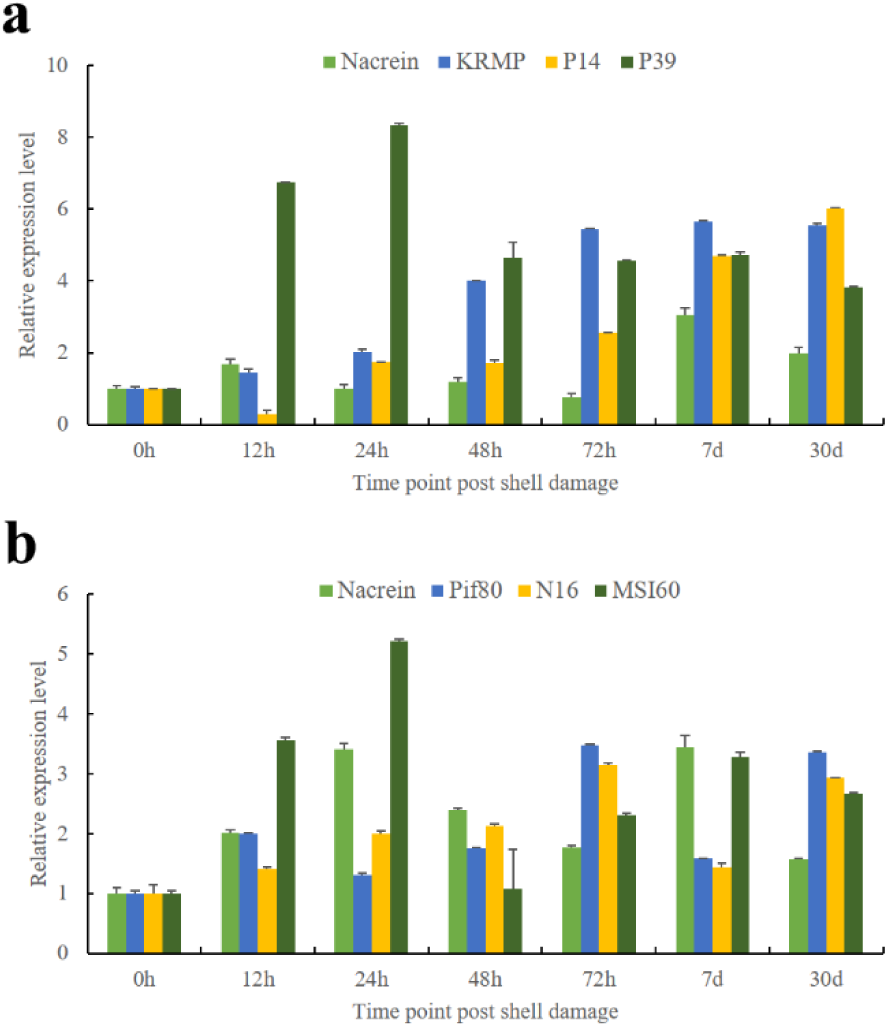
Gene expression of the shell matrix proteins in the mantle edge (a) and pallial zone (b) post artificial shell damage. h, hour; d, day. P14, Prismalin-14; P39, Prisilkin-39.

## 4. Conclusion

Shell formation of the pearl oyster *P. fucata* is mainly controlled by the mantle tissue. During the shell regeneration process, the shell damage, either artificial in the present study or natural in the open seawater, will cause the mantle tissue to retract and accelerate the secretion of SMPs. The retracted mantle tissue deposited an unusual prismatic layer upon the mature nacre sheet, and the upregulated SMPs promoted the CaCO_3_ nucleation. In this way the shell was quickly repaired, preventing secondly injury such as bacterial infection. However, how the physical signal of the shell damage is transferred to the mantle epithelial cells remains to be elucidated. Further study into the signal transduction pathway will shed light on the molecular mechanism underlying the precise regulation in the shell regeneration and eventually the shell mineralization in bivalves.

## Acknowledgements

This work was supported by National Natural Science Foundation of China Grants 31572594 and 31872543.

## Authors’ contributions

Jingliang Huang carried out the lab work, participated in data analysis, contributed to the design of the study and drafted the manuscript. Yangjia Liu and Taifeng Jiang coordinated the study. Wentao Dong and Guilan Zheng assisted in the data analysis. Liping Xie and Rongqing Zhang provided financial supports and revised the manuscript. All authors gave final approval for publication.

## Ethics statement

We confirm that the present study was approved by the Animal Ethics Committee of Tsinghua University, Beijing, China. All experiments were performed in accordance with relevant guidelines and regulations.

